# Segmental interpolation surface: a tool to dissect environmental effects on plant water-use efficiency in drought prone scenarios

**DOI:** 10.1101/033308

**Authors:** Gaston Quero, Sebastian Simondi, Omar Borsani

## Abstract

Water use efficiency (WUE), defined as the ratio of biomass (B) produced per unit of water transpired (E) by plants, is an important determinant of plant productivity. A mathematical approach was developed with the purpose explaining the WUE of two forage legumes (clover and birdsfoot trefoil). The approach applied involved an interpolation method by cubic spline which results in the smoothest curve that fits the set of data points obtained experimentally. The results obtained show the importance of recognizing the WUE as a function of two variables, one represents the supply (θ) and the other represents the demand (D). It is important to note that these surfaces generated by the model allowed the estimation of the WUE value for any value of θ and any value of D, showing it is able to dissect the effect of both environmental variables on WUE. From the surface generated by the model, a scalar field plot (SFP) was created. Analysis of these SFPs allowed decomposing the environmental effect on B and E parameters defining WUE. The SFPs allow identify, in one picture, what are the environmental conditions and what variable are explaining a high WUE in both species. Spline application for generating SFP could have a significant impact on the quantitative understanding of the WUE and our study represents a first step towards an analytical and integrated view of this parameters.

**Mathematics Subject Classification** 92B05 65D05 65D07 65D10

## 1. Introduction

Drought is one of the most important environmental factor that impact on plant and crop yield. Water soil availability is the single most important problem in global agricultural production (Rijsberman 2006) it is one of the main abiotic factors limiting crop production: losses in crop yield due to water stress exceed losses due to all other biotic and environmental factors combined (Lambers et al. 2008).

Relationship between the amount of water transpired by plants and dry matter accumulated thereof has been named in several ways: coefficient of transpiration, transpiration rate and plant water requirement, while the inverse of this relationship has been originally called transpiration efficiency, which more recently renamed as water use efficiency (WUE) (Jones 2004). The WUE reflects the balance between photosynthesis and transpiration.

WUE can be analysed at plant level, in a temporal scale of days and under environmental controlled conditions. Possibility to block the substrate evaporation by different methods reduces drastically the water loss by that way. Under the conditions describe above the WUE can be estimated as:

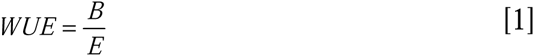
wherein B is aboveground biomass (g) and E the transpiration (Kg).

WUE quantified by this methodology analysis of genotype vs genotype can be performed. For instance if two genotypes (genotype *a* vs genotype *b*) are compared, applying equation [1] WUE of genotype *<u>a</u>* could be higher than the WUE of genotype *b* if the follow situation are resulting:

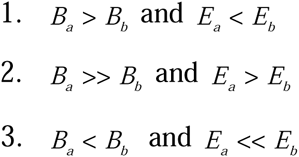

In situation 1 genotype *a* has more biomass than genotype *b* with less transpiration, in situation 2 genotype *a* has much higher biomass than genotype *b* with more transpiration. Finally, situation 3 shows how genotype *a* reduce the yield respect to genotype *b*, but with a higher reduction of transpiration respect to genotype *b*. Is important to note that the physiological behavior of the genotypes can be quite different in the three situations. Under drought conditions adaptive responses are triggered and reduction of E is part of these responses. Crop yield is a function of E and issue for the breeder because they want to reach genotypes with low E under stress with a minimum reduction in plant production (Blum 2005).

The capacity to develop an integrated point of view would be the basis to construct models of WUE which can be used under unpredictable environmental conditions and anticipate the consequences of drought on crop plants. So, modelling WUE represents a major objective and challenge for the scientific community with the final proposes of design more efficient and water-saving cropping systems (Damour et al. 2010).

Moreover, the complexity of the experimental observations requires an accurate analysis, and the need for a quantitative understanding of biological phenomena has led to analytical mathematical models. The study proposed here deals with this kind of models, i.e., reaching biologically meaningful results from mathematical models based on a simplification of experimental observations.

Cubic splines are widely used to fit a smooth continuous function through discrete data with piecewise cubic polynomials. The use of low-order polynomials is especially attractive for curve fitting because they reduce the computational requirements and numerical instabilities that arise with higher degree curves. These instabilities cause undesirable oscillations when several points are joined in a common curve. Cubic polynomials are most commonly used because no lower-degree polynomial allows a curve to pass through two specified endpoints with specified derivatives at each endpoint. The most compelling reason for their use, though, is that they guarantee continuous first and second derivatives across all polynomial segments. They have been applied to biological modelling at different levels of organizations from cellular or enzymatic processes (Zhan and Yeung 2011) to the parameterization of climate normal in large and heterogeneous regions (Rehfeldt 2006). In the area of crop science segmental splines have been used to describe the water soil characteristics (Kastanek and Nielsen 2001) and the nutrients dynamic flow from the soil to the plant (Claassen and Barber 1974).

However, segmented spline interpolations, to our knowledge, have never been applied to the analysis of WUE. Most empirical approaches use regression to determine a simple curve that “fits” the data, chiefly a relationship between WUE and air vapor pressure deficit (D). (Tanner, 1981; Kemanian et al. 2005; Damour et al. 2010). While there is a theoretical basis for the putative response of WUE to D, it is assumed that variation in the data is of lesser importance or simply due to experimental errors. In this context, the use of segmental spline interpolation appears as a new tool in modeling the WUE.

Unlike earlier approaches, a spline model respects each measurement and can provide greater sensitivity to unravel the factors that control WUE. This study tries to develop and applicate the segmental spline interpolation, in order to explain the environmental effect on WUE of two forage legumes in response to drought prone scenarios. Our medium-long term expectation is to develop a model able guide the improve WUE in crops through the identification of plant idiotype under a specific environment.

## 2. Materials, methods and algorithm

### 2.1 Plant material and growth conditions

*Lotus corniculatus* cv. San Gabriel (birdsfoot trefoil) and *Trifolium pratense* cv. LE116 (clover) seeds were obtained from M. Rebuffo INIA La Estanzuela (Colonia, Uruguay). The seeds were surface sterilized as described by Signorelli et al. (2013) and germinated at 28 °C for 2 days. Seedlings were transferred to 0.3 l pots containing a mix of river sand:vermiculite (1:1). The plants were grown under controlled conditions, 16 h light: 8 h dark cycle at 24/18 °C with a photosynthetic photon flux density of 250 μmol m^−2^ s^−1^. Plants were irrigated daily with Hornum’s nutrient solution (Handberg and Stougaard 1992). Eighth seedlings were planted in each pot and thinned to five seedlings after implantation.

### 2.2 Experimental design and treatments

Assays were performed in three independent growth chambers to generate three levels of air vapor pressure deficit (D) 0.94, 1.75 and 2.17 kPa were tested (Table 1). In each chamber the plants were grown for 30 days without water limitations, a time after which the pots were irrigated as to generate five water supply (θ) levels: 0.10, 0.15, 0.20, 0.25 and 0.30. Pots substrate surface was covered in order to minimize the water loss by evaporation of the substrate.

The θ were applied in each chamber in a complete randomized design with four replications per treatment. 30 days after transplanting four pots of each species were randomly sampled to record the initial plant weight. After 15 days in each substrate water content the remaining plants were harvested, dried to constant mass at 60 ºC and the final dry mass recorded. During the whole period of the experiment daily transpiration of each genotype in each θ were recorded.

**Table 1:**
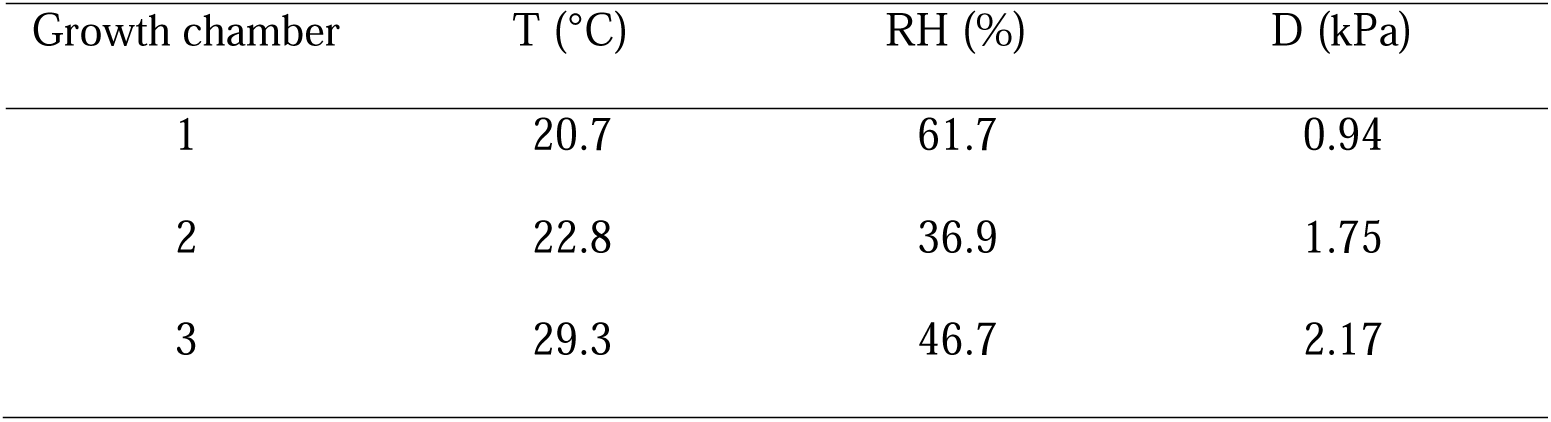
Atmospheric conditions in the three growth chambers

### 2.3 Water supply release curve and evaporative demand

The θ was determined gravimetrically by drying a subsample at 105 ºC for 72 h, and was calculated using the formula:

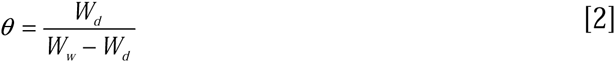

Where *W_w_* and *W_d_* were the wet and dry weight mass of substrates samples.

The D (kPa) were estimated according to the Tetens equation (1930) based on temperature and relative humidity:

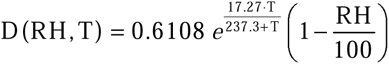

Wherein:

T is the room temperature in ºC and RH is the relative humidity in %

### 2.4 Algorithms

#### Polynomial interpolation

When there are two points, they can be joined by a straight line, i.e., determine a unique function of the first degree polynomial that passes through them. If we now have *n* points, a generalization of the above idea is to construct a polynomial of degree *n* that passes through them. It is a function of the form 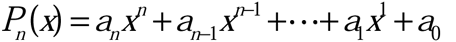 where *a*_0_,…,*a_n_* are real constants. The main reason for the use of the polynomial interpolation is that given any continuous function on a closed interval, there is a polynomial that is so close to the function as desired, this result is expressed precisely in the following theorem, known as Weierstrass Approximation Theorem; if *f* (*x*) is a continuous real-valued function defined on the real interval [*a*,*b*]. For every*ε* > 0, there exists a polynomial *P*(*x*) such that for all *x* in [*a*,*b*], we have |*f*(*x*)−*P*(*x*)| < *ε*.

While the construction of a single polynomial interpolation is justified theoretically, it presents many problems when the number of points to be interpolated is large. Firstly, the high degree polynomial functions tend to have big swings between interpolation points, which do not respond to phenomena being modeled and secondly its calculation is very complicated, which limits its usefulness in numerical analysis.

#### Segmental Polynomial Interpolation

An alternative method involves the split of the measurement interval into smaller subintervals, and using in each subinterval polynomials relatively low grade, trying to set the function bits thus has proper final appearance of the phenomenon being modeled. This type of interpolation is called segmental or spline, the most useful and widely used is that which is performed by polynomial degree three or cubic spline defined as follows:

Given a function *f* (*x*) defined as an closed interval [*a*,*b*] and a set of nodes *a*=*x*_0_ < *x*_1_ < … < *x_n_* = *b* L, a cubic spline interpolant *S*(*x*) for *f* (*x*) is a function that satisfies the following conditions:

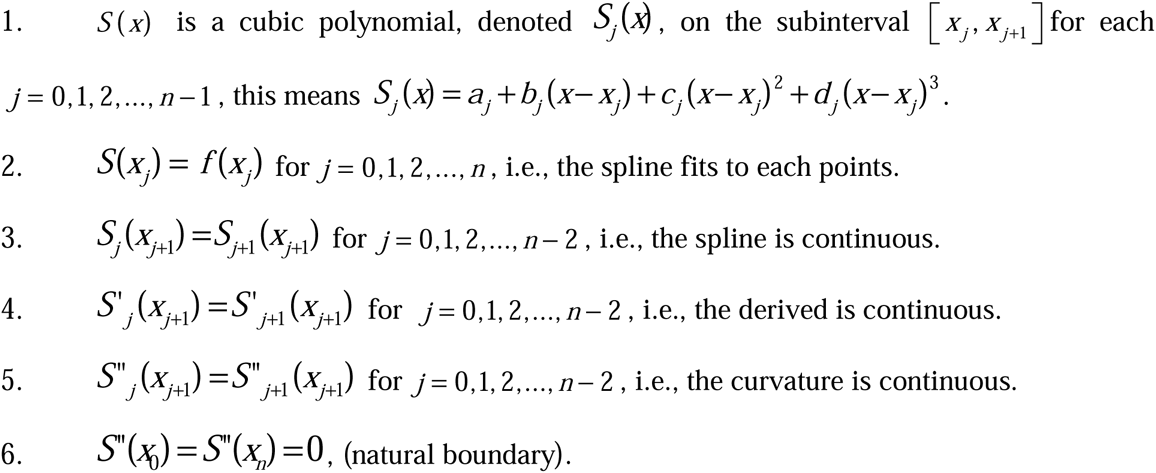

Under these conditions, the graph *S*(*x*) approximates the shape that a long flexible rod would assume if forced to go through the data points 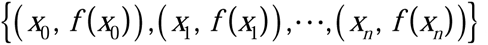, thereby minimizing the overall curvature of the graph.

#### Segmental interpolation surfaces

Using cubic splines to interpolating surfaces to build points 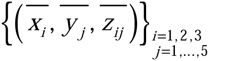 that contain the information of the data obtained experimentally as follows, the variable *x* represents any of the parameters of each test average demand (D, RH,T), *y* represents the supply parameter (θ) and the variable *z* represents the WUE, in the assay *i* under a matrix potential *j*. Each start intervals 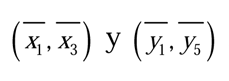 in fifty equal subintervals each, with the ends we build the next grid 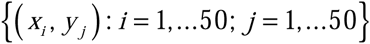 that will be nodes used to perform the interpolations. For each assay a cubic spline interpolant was constructed, the points 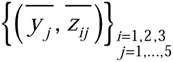 which are called 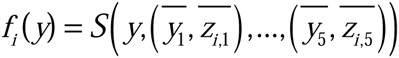. The points 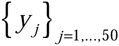 are valued on the grid in each spline generated, rending new nodes 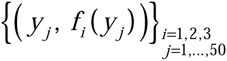. For each *j* value a cubic spline interpolant was constructed 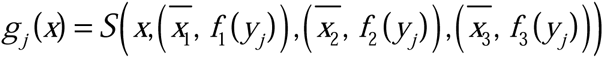.

The points 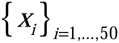 were valuated and the triples were armed 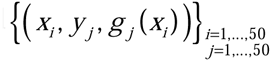 and with this point an interpolating surface were developed as is show in Results section.

## 3. Results

### 3.1 Experimental data

Figure 1 shows average values and deviations of WUE and E and B experimental obtained data from assay performed under three D and five θ conditions with the two legumes species. Figures 1A and 1B show that average values of WUE reached with D of 0.94 kPa were higher respect to the other D values in all θ. Also, with D values of 1.75 and 2.17 kPa no significantly differences were observed in that parameter. Figures 1C and 1D show E average values for each θ analyzed, for both species the E values with a D of 0.94 and 2.17 kPa were similar for all θ. A D of 1.75 kPa with a θ of 0.15 was the only conditions which generated a significantly lower E values. Figures 1E and 1F show that the average values of biomass for each specie in all θ are higher with a D of 0.94 kPa respect to the other two assay. In spite of the means values of water transpirated in each θ for the assay at D of 0.94 and 2.17 kPa being similar, the biomass produced at the lower D was always the highest in all θ. This implies that necessarily the higher WUE was reached with lower water demand.

**Figure 1.**
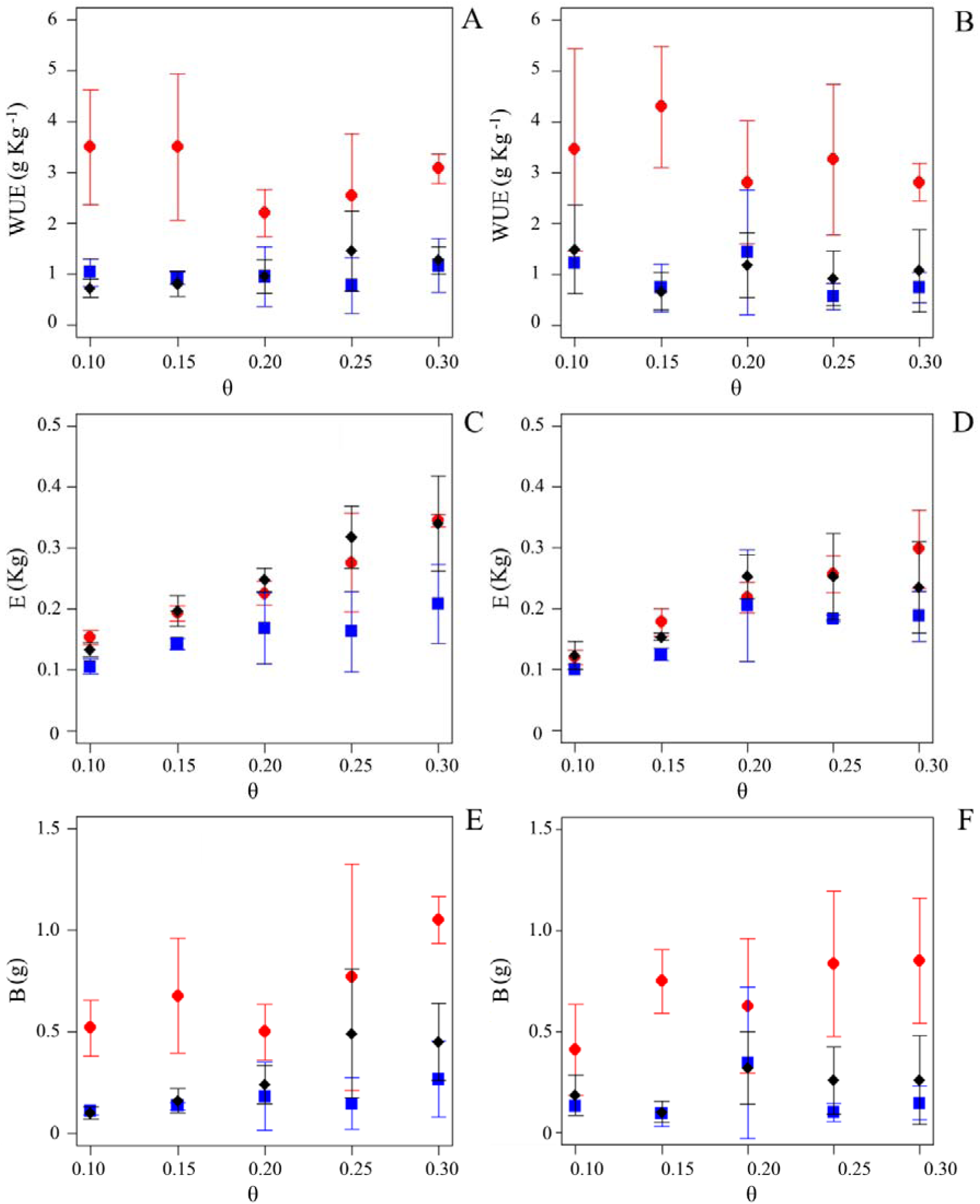
Experimental data obtained from assays performed under three air vapor pressure deficit (D) and five water supply (θ). A and B Water use efficiency (WUE) of birdsfoot trefoil and clover respectively. C and D Transpiration (E) of birdsfoot trefoil and clover respectively. D and F Biomass yield (B) of birdsfoot trefoil and clover respectively. Results are expressed by means and deviation. Red circles represent the results obtained from conditions generated with a D of 0.94 kPa, blue squares from conditions generated with a D of 1.75 kPa and black diamonds from conditions generated with a D of 2.17 kPa.

### 3.2 Inputting the model with experimental data

Application of segmental spline interpolation to the average of the WUE exposed in the figure 1A and 1B, resulted in the curves shown in Figure 2A and 2B (red curve at D of 0.94 kPa; blue at D of 1.75 kPa, and black at D of 2.17 kPa). These curves show that the results of the assay made at D of 0.94 kPa are significantly higher along the range of θ respect to those obtained in the other two assays for both species, as we are observed as in Figure 1.

In the figure 2A is observed how in birdsfoot trefoil the WUE curves generated by the model, at D of 1.75 y 2.17 kPa (blue and black respectively) have inverted increase and decrease intervals. For clover (figure 2B), the WUE at D of 0.94 kPa has inverted increase and decrease intervals when was compared with the simulation generated for the other two assays. For each legume specie, we used the curves plotted in Figure 2A and 2B to evaluate the fifty nodes (see section 2 M&M), in which the range of θ was divided, obtaining values from which we construct the interpolating surfaces by adding a new dimension, in this case the D. Figures 2C and 2D are the surfaces of WUE relative to D and θ for birdsfoot trefoil and clover respectively.

Surfaces show a direct dependence of WUE of D and θ. In both species at low D values the WUE is highly dependent of θ, but this dependency decreases when D increases. It is important to note that these surfaces allow us to estimate the WUE value for any value of θ between 0.10 and 0.30 and any value of D between 0.94 and 2.17 kPa. For instance, the cross section of surfaces in 2C and D allow the generation of curves which modelling the WUE as function of D for each θ analyzed (Figures 2E y F).

**Figure 2.**
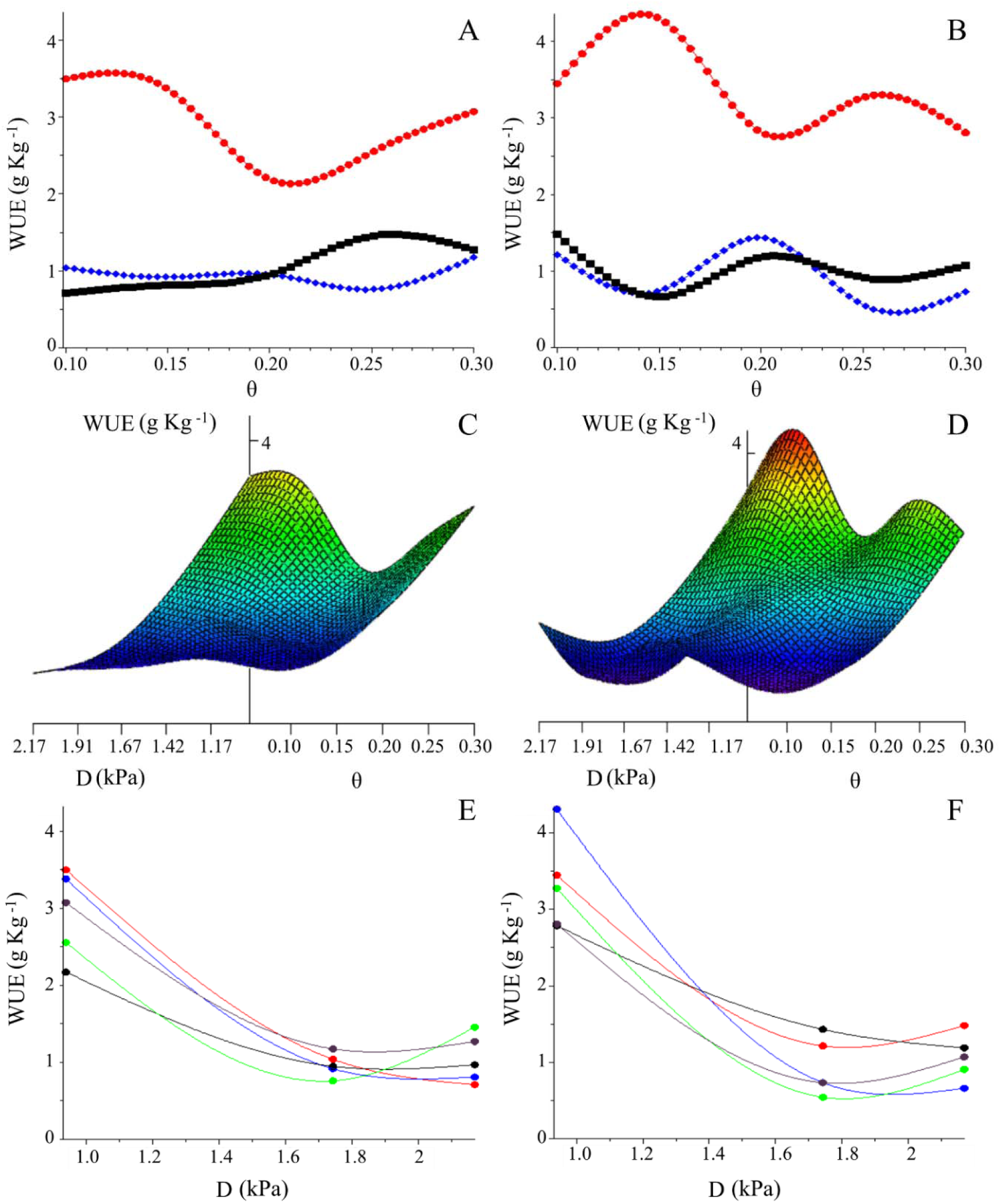
Water Use Efficiency (WUE) as function of air vapor pressure deficit (D) under five supply (θ) conditions. A and B Spline curves interpolate the average values of the WUE of birdsfoot trefoil and clover respectively. Red curve indicate interpolation of data obtained at D of 0.94 kPa; blue at D of 1.75 kPa, and black at D of 2.17 kPa. C and D WUE surfaces depending of D and θ para birdsfoot trefoil and clover respectively. E and F cross sections of Figures 2C (birdsfoot trefoil) and 2D (clover), centered on the average experimental θ data. Red curve was generated from data at θ of 0.10, blue curve at 0.15, black curve at 0.20, green curve at 0.25 and violet curve at 0.30.

### 3.3 Differential legume species responses to D and θ

WUE defines an index resulting from the ratio between the B produced and the E. Figure 3A shows the overlapping of the surfaces 2C and 2D and reveal wherein one species is more efficient in the WUE than the other.

In Figure 3B, in blue is shown the D and θ coordinates where the clover has a higher WUE than birdsfoot trefoil and in red ones where this ratio is inverted. In analogy, in the figures 3C and 3D the blues points show the coordinates where clover would have higher E and B respectively, compared with birdsfoot trefoil.

**Figure 3.**
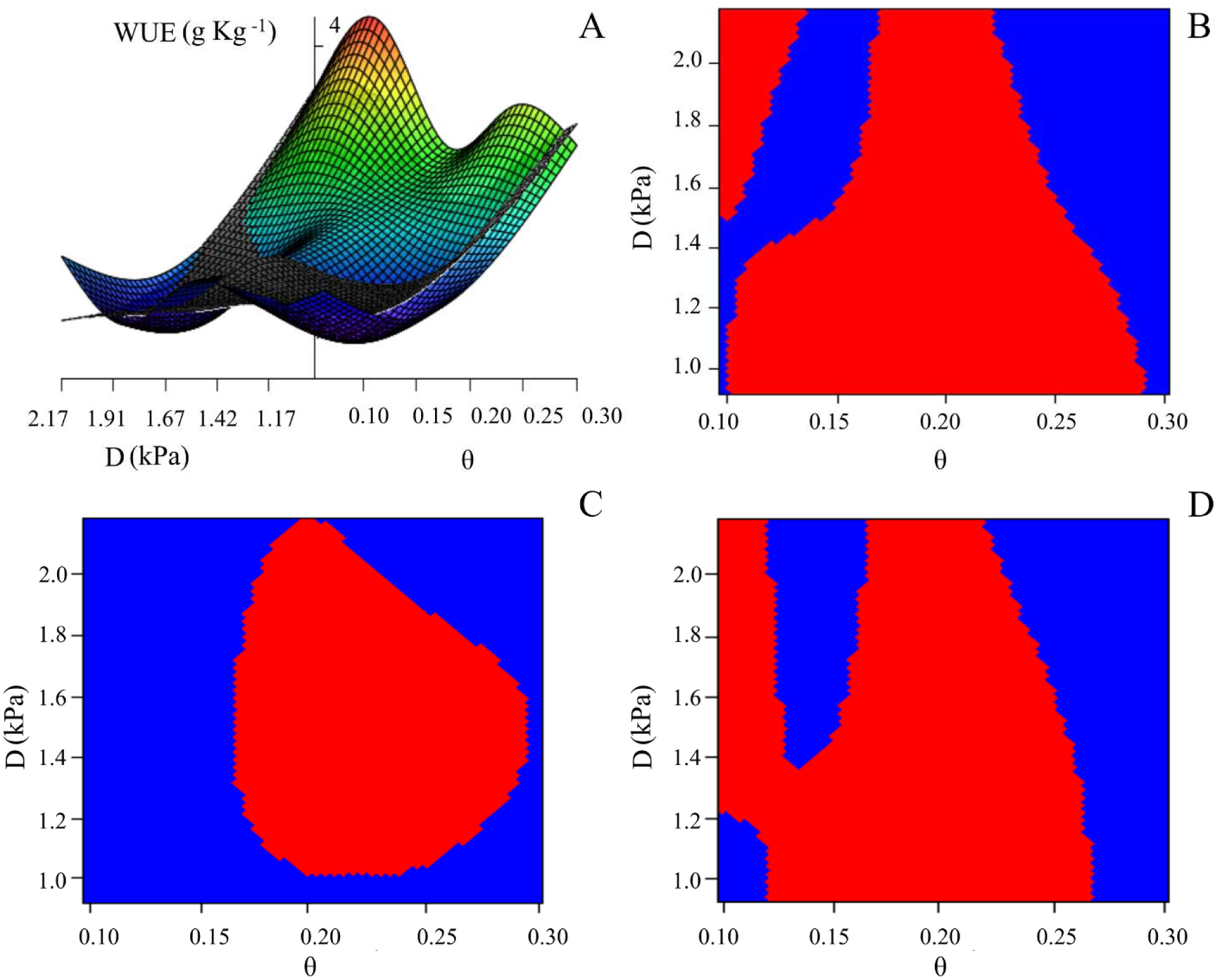
Surface and Scalar Field Plots (SFPs) of Water Use Efficiency (WUE), Transpiration and Biomass as function of air vapor pressure deficit (D) under five water supply (θ) conditions. A. WUE overlapped surfaces of the birdsfoot trefoil and clover. B, C and D, SFPs of WUE, transpiration and biomass yield of birdsfoot trefoil and clover respectively. In this figures the blue points indicate conditions when the clover is higher of birdsfoot trefoil and the red points when this ratio is inverted.

**Figure 4.**
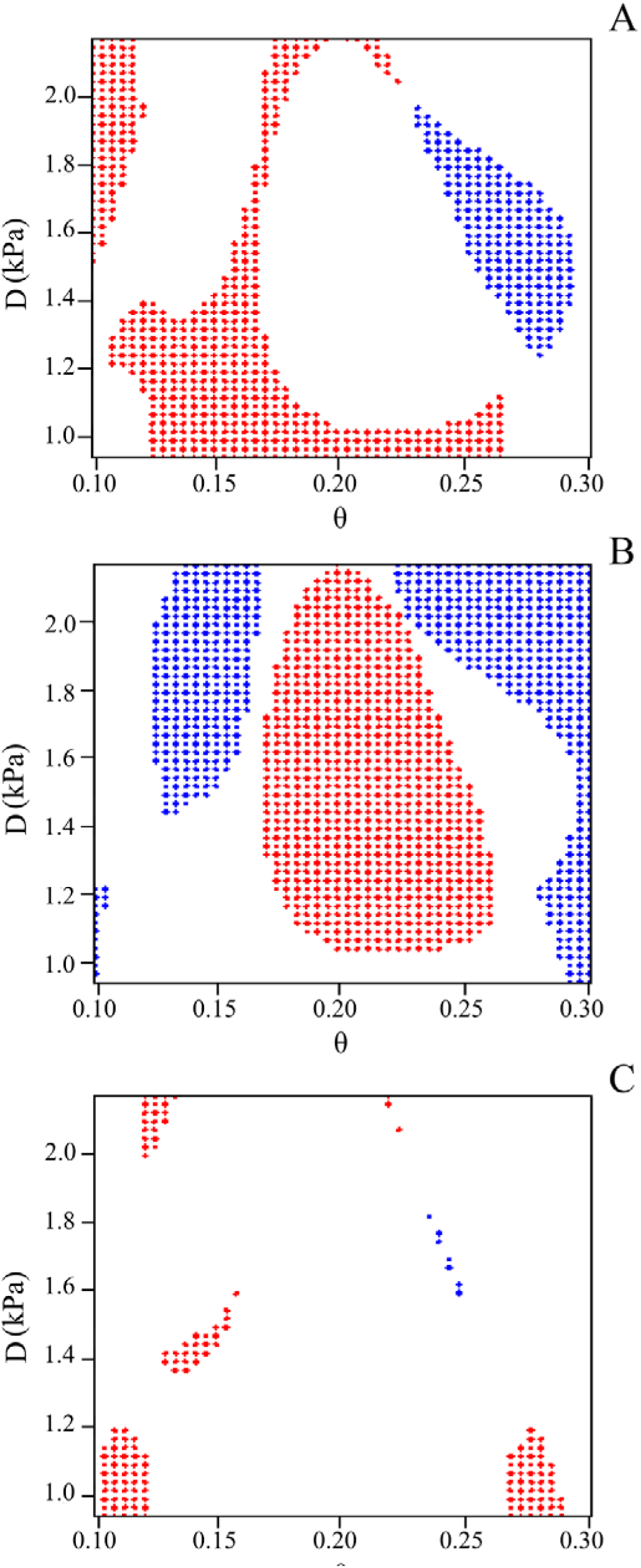
Dissection of WUE components as function of air vapor deficit (D) and water supply (θ). A. WUE generated by a high biomass production (B) with low transpiration (E). B. WUE generated by a much higher B but with a high E. C. WUE generated by a low B but with a much lower E. Blue and red points indicate conditions when the highest WUE correspond to clover and birdsfoot trefoil respectively.

## 4. Discussion

Several efforts are being carried out in order to improve water deficit tolerance in crops, however the complexity of the responses involved and the strong interaction of genotype with the environment has delayed the identification of promising ways to reach the selected goal (Malosetti et al. 2013). One physiological parameter that has been included in plant phenotyping to improve drought performance in crops is the water use efficiency (WUE). This parameter results from the behaviour of several physiological variables (conductance, photosynthesis, and respiration) so it could be a good candidate to be applied with a crop breeding goal. However, as is the case with many other plant physiological parameters, it is highly influenced by environmental variables like air vapour pressure deficit, relative humidity, temperature, soil water content and the interaction among them. In this way, modelling and simulation of WUE changes under different environmental condition becomes a critic requirement for its use as phenotypic marker.

The use of segmental spline for modelling the WUE is primarily due to two of its major properties. First, it does not have error adjustment, since the generated surface passes through all the average values obtained from the WUE assays. Second, it has the property of minimum curvature, i.e., it provides “softer” average surface interpolating points (Mathews et al. 2000). Another way to understand this property is that the cubic spline interpolation minimizes the potential energy that is the strain energy that makes minimal oscillatory behaviour. This could explain why a purely mathematical model obtains the same relations between the WUE and the D than the functional models (Kemanian et al. 2005; Wang et al. 2007). These relationships are obtained performing cross sections to interpolating surfaces of WUE generated as a function of the θ and the D, at fixed θ (Figure 2E and 2F), thus showing that the model presented here is a generalization of functional models.

The model can also be used as an attractive tool for massive plant phenotyping strategies; because it allows evaluation in contrasting conditions. Since from few quantities of data it is possible to estimate the WUE over the range of measurements taken with differential response of the genotype, which is also perceived by the model. The potential of this model is capturing the key principles governing the phenomenon of WUE. A quantitative understanding of these principles leads to new information that has not been discovered by experimental methods. Successful models of mathematical analysis lead to insights that are unreachable experimentally.

The analytical model presented allows estimating responses, inferring functionality and designing different experimental scenarios for future assays; it can also be adjusted and updated with new data. The results obtained show the importance of recognizing the WUE as a function of two variables, one represents the supply (θ) and the other represents the demand (D).

WUE overlapping surfaces generated as a function of θ and D (figure 2C and 2D) are shown in figure 3A. From which we generate a scalar field plot (SFP) (figure 3B) where the coordinates of D and θ are shown in blue in which the clover has a higher WUE and in red birdsfoot trefoil where this relationship is reversed. By proceeding in the same way, we generate the SFPs comparatives for transpiration (figure 3C) and biomass yield (figure 3D). Taking these three pictures is possible to determine what relationship between biomass and transpiration resulting in a high WUE.

The analytical model applied here allows the analysis of the results of the SFPs and identifying in one picture what are the environmental conditions and what variables are explaining a high WUE in both species. For instance, once the zones are defined the model is able to identify the coordinates where it is possible to find a high WUE generated by a high biomass yield with a high transpiration rate (Figure 4B). On the other hand, it is possible to define coordinates to obtain high WUE with low biomass yield conditioned by a high reduction of transpiration rate (Figure 4C). However, the most outstanding propriety of this model is the capacity to identify coordinates wherein the WUE generates a high biomass yield with a low transpiration rate (Figure 4A). This region could identify genotypes with promising traits, so this analysis could potentially be used as genotype selection methods.

Application of this kind of mathematical modelling methodology is an attractive way to contribute to the study of complex parameter as WUE. Future analysis with massive data input will define if this approach appears as new tool for the crop breeding programs.

## Acknowledgements

This work was supported by Programa Grupo I+D 418 Comisión Sectorial Investigación Científica-Universidad de la Republica (GQ and OB) and Secretaria de Ciencia Técnica y Posgrado-Universidad Nacional de Cuyo M019 (SS).

